# Large Language Model Agent for Modular Task Execution in Drug Discovery

**DOI:** 10.1101/2025.07.02.662875

**Authors:** Janghoon Ock, Radheesh Sharma Meda, Srivathsan Badrinarayanan, Neha S Aluru, Achuth Chandrasekhar, Amir Barati Farimani

## Abstract

We present a modular framework powered by large language models (LLMs) that automates and streamlines key tasks across the early-stage computational drug discovery pipeline. By combining LLM reasoning with domain-specific tools, the framework performs biomedical data retrieval, domain-specific question answering, molecular generation, property prediction, property-aware molecular refinement, and 3D protein–ligand structure generation. In a case study targeting BCL-2 in lymphocytic leukemia, the agent autonomously retrieved relevant biomolecular information—including FASTA sequences, SMILES representations, and literature—and answered mechanistic questions with improved contextual accuracy over standard LLMs. It then generated chemically diverse seed molecules and predicted 67 ADMET-related properties, which guided iterative molecular refinement. Across two refinement rounds, the number of molecules with QED > 0.6 increased from 34 to 55, and those passing at least four out of five empirical drug-likeness rules rose from 29 to 52, within a pool of 194 molecules. The framework also employed Boltz-2 to generate 3D protein–ligand complexes and provide rapid binding affinity estimates for candidate compounds. These results demonstrate that the approach effectively supports molecular screening, prioritization, and structure evaluation. Its modular design enables flexible integration of evolving tools and models, providing a scalable foundation for AI-assisted therapeutic discovery.

## Introduction

The discovery of drug-like molecules for treating specific diseases is fundamental to therapeutic innovation and the advancement of public health. However, identifying viable candidates is a highly complex and resource-intensive process. It requires navigating an enormous chemical space, integrating heterogeneous biological and chemical data sources, and conducting iterative experimental validation. Collectively, these challenges contribute to drug development timelines of 10–15 years and costs exceeding $2 billion per approved therapy.^1–3^

To mitigate these challenges, computational approaches have become essential in early-stage drug development. Methods such as molecular docking,^4–6^ quantitative structure–activity relationship (QSAR) modeling,^7–9^ and molecular dynamics simulations ^10,11^ have significantly improved the speed and accuracy of compound screening and optimization. More recently, machine learning (ML) techniques have expanded this toolkit by enabling predictive modeling of pharmacokinetic and pharmacodynamic properties, de novo molecular generation, and multitask learning across diverse biomedical tasks. ^12–15^ Notably, AlphaFold has achieved high-accuracy protein structure prediction,^16–18^ while Graph Neural Networks (GNNs) and transformer-based models improve property prediction.^19–22^ In parallel, generative models such as Variational Autoencoders (VAEs), Generative Adversarial Networks (GANs), and diffusion models (e.g., JT-VAE, MolGAN, MolDiff) enable de novo or target-specific molecular design.^23–25^ Collectively, these methods increase the scalability and automation potential of modern drug discovery workflows.

Despite this progress, most computational methods remain task-specific and require manual orchestration by domain experts. However, drug discovery is inherently a multi-step, interdependent process that requires seamless integration across diverse tasks.^1–3^ This fragmented implementation limits the scalability and efficiency of current pipelines. As the demand for faster and more cost-effective drug screening grows, there is an urgent need for unified, intelligent platforms capable of autonomously coordinating these tasks while supporting expert decision-making.

Large Language Models (LLMs) offer a compelling solution to this need. Trained on massive corpora of natural language data, LLMs exhibit strong reasoning capabilities and domain-agnostic knowledge. When augmented with external tools, such as domain-specific plugins, APIs, and software libraries, LLM-based agents can overcome the limitations of general-purpose language models and act as interpretable, flexible controllers of scientific workflows.^26^ LLM agents have recently been applied to automate diverse aspects of scientific discovery, including experimental design, material synthesis planning, and data analysis.^27–31^ For instance, Boiko et al. introduced Coscientist, an LLM agent capable of autonomously planning and executing chemistry experiments, significantly enhancing lab productivity while reducing human intervention.^32^ Similarly, the dZiner framework leverages LLM agents for molecular design through iterative reasoning and structure-based optimization.^33^

In this work, we introduce AgentD, an LLM-powered agent framework designed to support and streamline the drug discovery pipeline. The agent performs a wide range of essential tasks, including biomedical data retrieval from structured databases and unstructured web sources, answering domain-specific scientific queries, generating seed molecule libraries via SMILES-based generative models, predicting a broad spectrum of drug-relevant properties, refining molecular representations to improve drug-likeness, and generating 3D molecular structures for downstream analysis. Our results demonstrate that this agent-driven framework streamlines early-phase drug discovery and serves as a flexible foundation for scalable, AI-assisted therapeutic development. Furthermore, its modular architecture allows for continual improvement as more advanced tools and models become available.

## Agent Design

### Task Modules

AgentD performs six essential tasks across the drug discovery pipeline, as illustrated in Figure 1. The agent leverages large language models from providers such as OpenAI and Anthropic; in this study, we primarily use OpenAI’s GPT-4o as both the central reasoning engine and the natural language interface. By integrating domain-specific tools and databases, AgentD coordinates a wide range of activities - from data retrieval and molecular generation to property evaluation and structure prediction. The primary tool components supporting each task module are summarized in Table 1.

**Figure 1:**
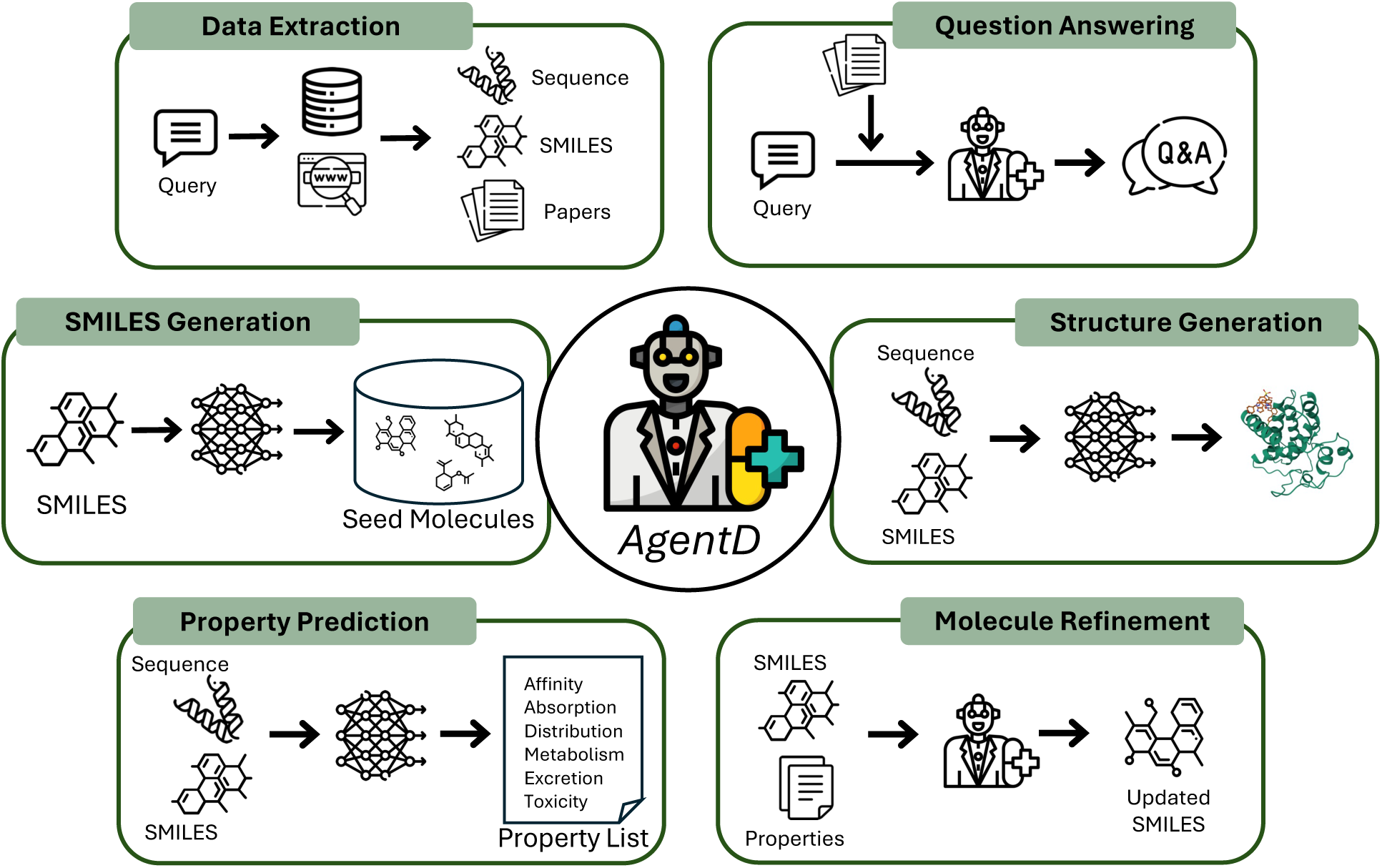
Overview of task modules supported by AgentD. In the question answering and molecule refinement tasks, outputs are generated directly by the language model, whereas in other tasks, the final results are produced by integrated external tools.

**Table 1:** Key external tools integrated into each task module within the AgentD drug discovery pipeline.

| Task Module | Primary Tool Components |
| --- | --- |
| Data Extraction | UniProt, ChEMBL, Serper, Semantic Scholar APIs |
| Question Answering | RAG |
| SMILES Generation | REINVENT, Mol2Mol |
| Property Prediction | DeepPK API, BAPULM |
| Molecule Refinement | LLM internal knowledge and reasoning<sup>*</sup> |
| Structure Generation | Boltz-2 |

<sup>\*</sup>Adapted from the dZiner framework.<sup>33</sup>

#### Data Extraction

AgentD is capable of retrieving biomedical information from both structured and unstructured sources. Given a query - such as identifying known drugs associated with a specific protein–disease context - the agent retrieves the protein’s FASTA sequence from the UniProt database,^34^ searches for relevant drug names via the web, and extracts corresponding SMILES representations from the ChEMBL database.^35^ Additionally, AgentD autonomously constructs keyword-based queries to download relevant open-access scientific literature from Semantic Scholar, providing a broader context for downstream reasoning and decision-making.

#### Question Answering

In therapeutic applications, high accuracy and mechanistic specificity are critical. Generic responses that only appear plausible are inadequate for addressing domain-specific questions relevant to drug discovery and biomedical research. To handle domain-specific scientific queries, AgentD employs a retrieval-augmented generation (RAG) strategy. By grounding its responses in literature obtained during the data extraction phase, the agent provides context-aware and evidence-based answers.

#### SMILES Generation

Constructing a diverse and chemically relevant seed molecule pool is essential for effective early-stage virtual screening. AgentD generates seed molecules in SMILES format using external generative models. In this study, we incorporate two such models: REINVENT,^36,37^ which supports de novo molecule generation without requiring input SMILES, and Mol2Mol,^38^ which performs conditional generation to produce molecules structurally similar to a given input SMILES. This dual capability enables both exploration and exploitation in the molecular search space.

#### Property Prediction

For each candidate molecule, AgentD predicts key pharmacologically relevant properties, including ADMET (absorption, distribution, metabolism, excretion, and toxicity) profiles and binding affinity (e.g., pKd). ADMET prediction is performed using the Deep-PK API,^39^ which accepts SMILES strings as input. For binding affinity estimation, the BAPULM model^40^ is used, which operates on both the SMILES and the protein’s FASTA sequence. These predictions help prioritize compounds based on both efficacy and safety.

#### Molecule Refinement

Based on the predicted properties, AgentD can identify molecular shortcomings such as toxicity or poor permeability, and propose targeted structural modifications to improve attributes like solubility and metabolic stability. This SMILES refinement is carried out solely through the LLM’s internal reasoning and built-in chemical knowledge, without additional model-based tools.^33^

#### Structure Generation

AgentD can generate 3D structures of protein–ligand complexes using Boltz-2 as an external tool.^41,42^ This process produces candidate complex structures along with associated binding metrics such as IC_50_ values and inhibitor probability. These structures can serve as inputs for downstream computational tasks such as docking simulations or molecular dynamics, offering deeper insight into the biophysical interactions of the drug candidates.

### Workflow

All six task modules in AgentD are designed to support key components of the drug discovery pipeline, as illustrated in Figure 2. In this study, we demonstrate the workflow using a case study involving BCL-2, a well-characterized therapeutic target in lymphocytic leukemia. The process begins with a user-provided query - for instance, identifying drug molecules that target a specific disease-related protein. Through the data extraction module, AgentD retrieves the protein’s FASTA sequence from UniProt and identifies known drugs using sources such as ChEMBL and Google search APIs.

**Figure 2:**
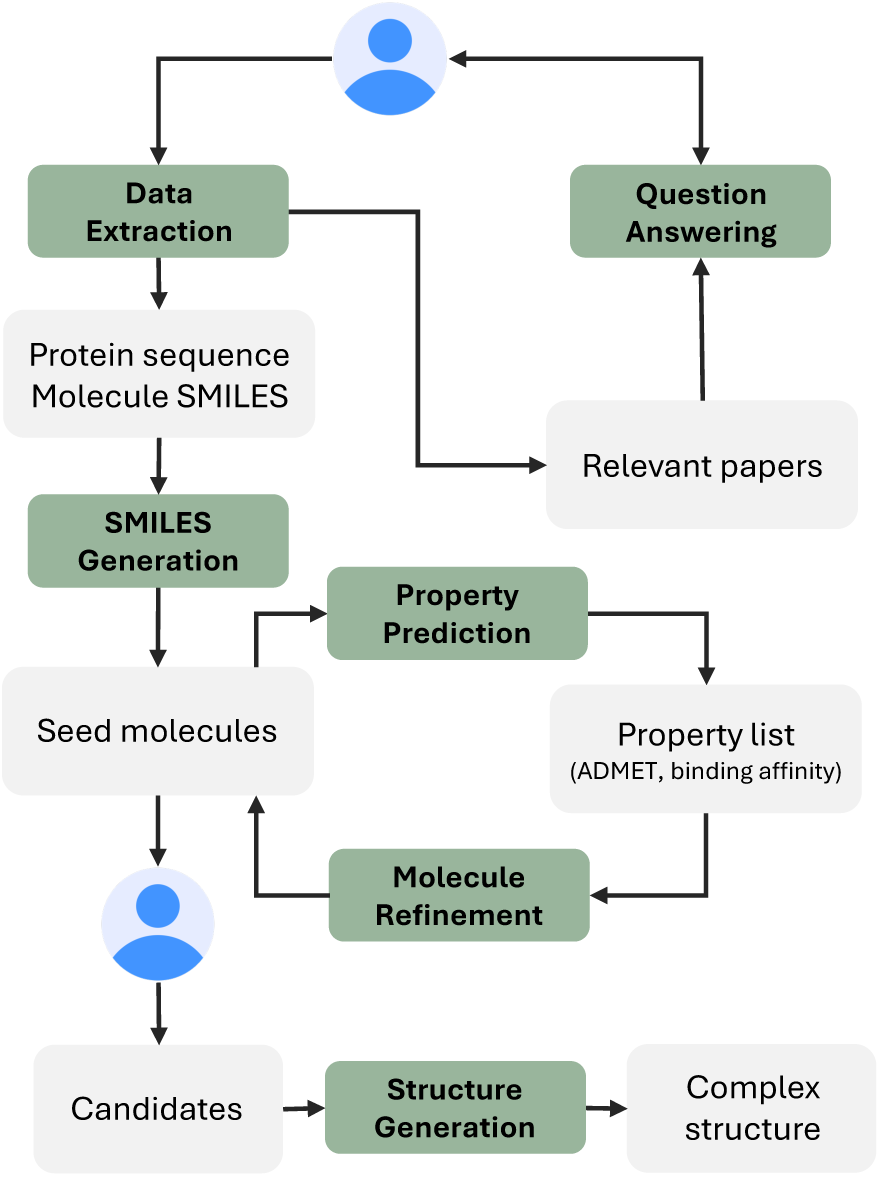
Workflow illustrating how each task module within AgentD supports the drug discovery pipeline.

The SMILES of a known drug serves as a starting point for the SMILES generation task, which builds a chemically diverse library of candidate molecules using generative models. These molecules are then evaluated using property prediction tools to estimate key pharmacological characteristics such as ADMET profiles and binding affinity. Based on these predicted properties, AgentD proposes SMILES-level modifications to improve attributes like solubility, toxicity, and metabolic stability. These refinements rely on the language model’s internal reasoning rather than external optimization tools and help enrich the candidate pool.

Following refinement, domain-specific criteria can be applied by the user to select promising compounds. For these, the structure generation task creates 3D protein–ligand complexes using the ligand SMILES and target protein sequence. This enables more detailed downstream analyses. The structure generation step also outputs auxiliary metrics like predicted IC_50_ and inhibitor probability as proxies for binding strength. While these estimates, derived from Boltz-2, offer useful guidance, they should be interpreted cautiously, as they are not highly accurate.

Throughout the workflow, users may request clarification or validation of scientific concepts. The question answering module, implemented using RAG, supports this by leveraging literature collected during the data extraction phase. This functionality serves both to enhance the pipeline and to address user-specified scientific queries on demand.

## Results

### Data Extraction

Reliable access to protein sequences and ligand structures is essential for computational tasks such as virtual screening, molecular modeling, and property prediction. To support this capability, AgentD integrates web search functionality and database API access as tool modules. When provided with a user query specifying a target protein and associated disease, the agent is instructed to: (i) retrieve the protein’s FASTA sequence, (ii) identify existing drugs relevant to the specified target–disease context, and (iii) download related open-access scientific literature.

The agent queries the UniProt database for the protein sequence and uses web search tools (via the Serper API) to identify relevant drug molecules. Once a drug is identified, its SMILES representation is retrieved from the ChEMBL database. In parallel, the agent generates keyword-based queries to download supporting publications from Semantic Scholar. To demonstrate this functionality, the agent was tested on example queries including BCL-2 for lymphocytic leukemia, JAK2 for myelofibrosis, and KIT for renal cell carcinoma. The results, summarized in Figure 3, show that the agent successfully retrieved key molecular data, such as identifying venetoclax as a BCL-2 inhibitor.

**Figure 3:**
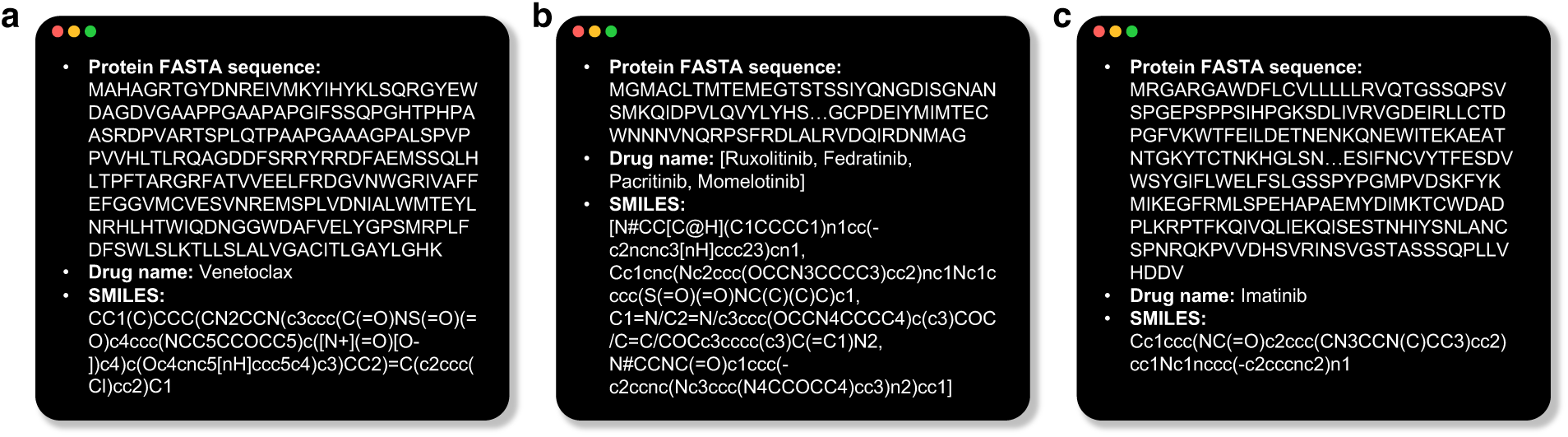
Examples of the data extraction. Given a user query specifying a target protein and associated disease, AgentD retrieves the protein’s FASTA sequence and identifies existing drug molecules related to the query.

The extracted information is used in several downstream modules. The retrieved SMILES is used to initialize molecule generation models for building a seed library. Both the SMILES and protein sequence serve as inputs to the property prediction and 3D structure generation components. Additionally, the downloaded documents are embedded into a vector database, enabling context-aware retrieval during the agent’s question answering task.

### Domain-Specific Question Answering

To address the limitations of generic, unsupported answers in biomedical contexts, AgentD employs RAG to ground its responses in domain-specific literature. During the data extraction phase, the agent formulates keyword-based queries from the target protein and disease, retrieves open-access papers via Semantic Scholar, and stores them in a vector database for later retrieval during question answering.

We evaluated this capability using a representative study by Weller et al., which describes how venetoclax activates the integrated stress response (ISR), leading to NOXA upregulation and MCL-1 inhibition.^43^ Based on this paper, we formulated domain-specific questions and compared the responses generated by AgentD with RAG to those from the standard GPT-4o model (Figure 4).

**Figure 4:**
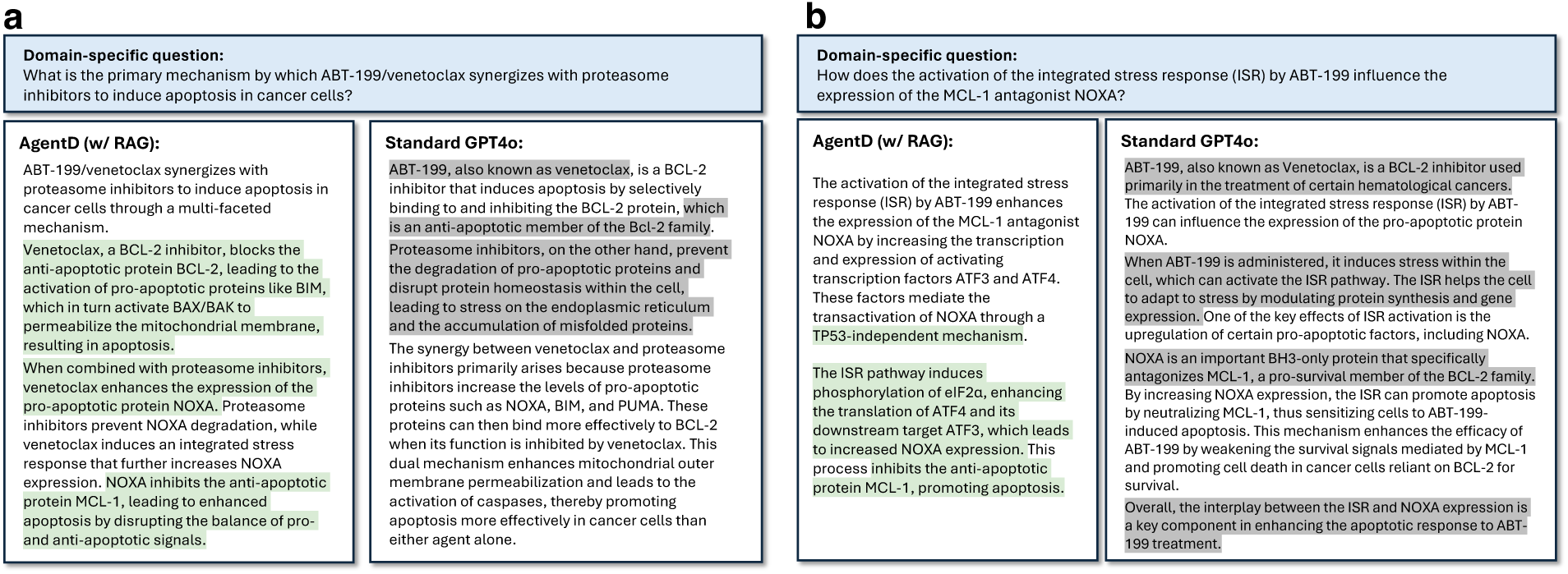
Comparison of question answering performance between AgentD with RAG and the standard GPT-4o model. Green highlights indicate grounded, context-relevant information supported by the reference paper, while grey highlights denote generic content not directly related to the question.

AgentD consistently generates more detailed and mechanistically accurate responses. For example, when asked about the synergy mechanism between venetoclax and proteasome inhibitors, the RAG-augmented answer includes key elements such as ISR activation, ATF3/ATF4-mediated transcription of NOXA, and downstream effects involving BIM, BAX/BAK activation, and mitochondrial membrane permeabilization—closely aligning with the mechanistic explanation presented in the source paper (Figure 4a). In contrast, GPT-4o provides a more generic explanation, omitting critical components such as the ISR pathway and transcriptional regulators.

A similar pattern is observed when asked how ABT-199 influences NOXA expression via ISR. AgentD correctly references eIF2*α* phosphorylation, TP53 independence, and the ATF3/ATF4 regulatory cascade. While GPT-4o’s response is fluent, it lacks these essential mechanistic details. As shown in Figure 4, AgentD’s context-aware answers are highlighted in green, whereas GPT-4o’s generic responses appear in grey.

### Seed Molecule Generation

The initial library of molecules serves as a critical foundation for exploring chemical space and identifying candidates for further optimization. To construct this seed molecule pool, AgentD leverages two complementary generation strategies. REINVENT enables de novo molecule generation, allowing for broad and unbiased exploration of chemical space without the need for an input structure. In contrast, Mol2Mol performs conditional generation, producing analogs that are structurally similar to a specified input molecule. In our workflow, the existing drug identified during the data extraction phase—such as venetoclax—is used as input for Mol2Mol, enabling the agent to focus on chemically relevant and biologically meaningful regions of the search space.

After retrieving the SMILES of the known drug, AgentD automatically integrates it into a configuration file and executes both REINVENT and Mol2Mol to generate the initial seed molecules. As shown in Figure 5, REINVENT-generated molecules are widely distributed across the chemical space, reflecting its exploratory capabilities, whereas Mol2Mol-generated molecules are more tightly clustered around the input molecule, enabling targeted exploration near known active scaffolds. This seed library serves as the starting point for downstream tasks such as property prediction, refinement, and structure generation.

**Figure 5:**
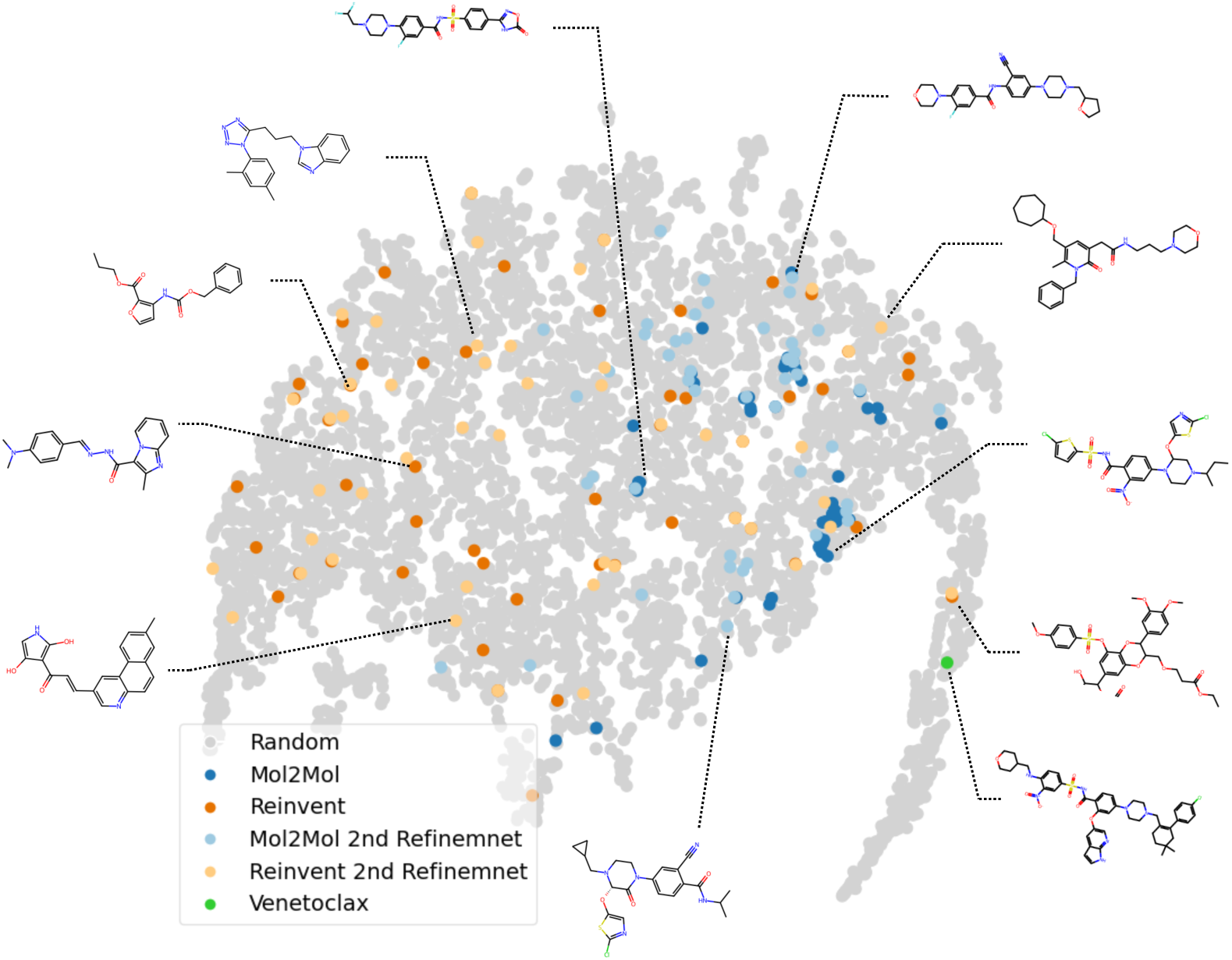
t-SNE visualization of chemical space, based on Mordred-derived molecular descriptors, ^44^ including seed molecules generated by AgentD (Mol2Mol, REINVENT), refined molecules after property-aware optimization, 5,000 randomly sampled compounds from ChEMBL, and the reference drug venetoclax.

### Property-Aware Molecular Refinement

To enhance the quality of the seed molecule pool, AgentD performs property-driven refinement of SMILES structures by identifying and addressing limitations that compromise their drug-likeness. Using the Deep-PK model API,^39^ the agent predicts 67 ADMET properties including absorption (e.g., Caco-2 log *P*_app_), distribution (e.g., blood–brain barrier permeability), metabolism (e.g., CYP1A2 inhibition), excretion (e.g., drug half-life), and toxicity (e.g., biodegradability, general toxicity, and logD), as well as 7 general physicochemical properties such as lipophilicity and solubility. These properties are critical for evaluating pharmacokinetic profiles and safety. A complete list of predicted ADMET properties is provided in Table S1. In parallel, AgentD uses the BAPULM model^40^ to estimate binding affinity (pKd) from the ligand SMILES and the protein FASTA sequence. Details of both models are described in the Methods section.

Based on these predictions, AgentD identifies unfavorable molecular properties relevant to drug development such as low permeability or high toxicity, and proposes targeted structural edits to improve them. A comprehensive list of risk-associated properties is provided in Table S2. We demonstrate this refinement process over two iterations, beginning with an initial set of 100 molecules. These iterations produced 99 and 95 additional valid SMILES, respectively, which were added to the candidate pool. To illustrate how the refinement works in practice, Figure 6a shows a molecule initially flagged for poor permeability due to a low predicted log *P*_app_ value. In response, AgentD suggested replacing a hydroxyl group with a methyl group in the first round, followed by substituting a sulfonamide with an amine in the second-both changes contributed to improved predicted permeability.

**Figure 6:**
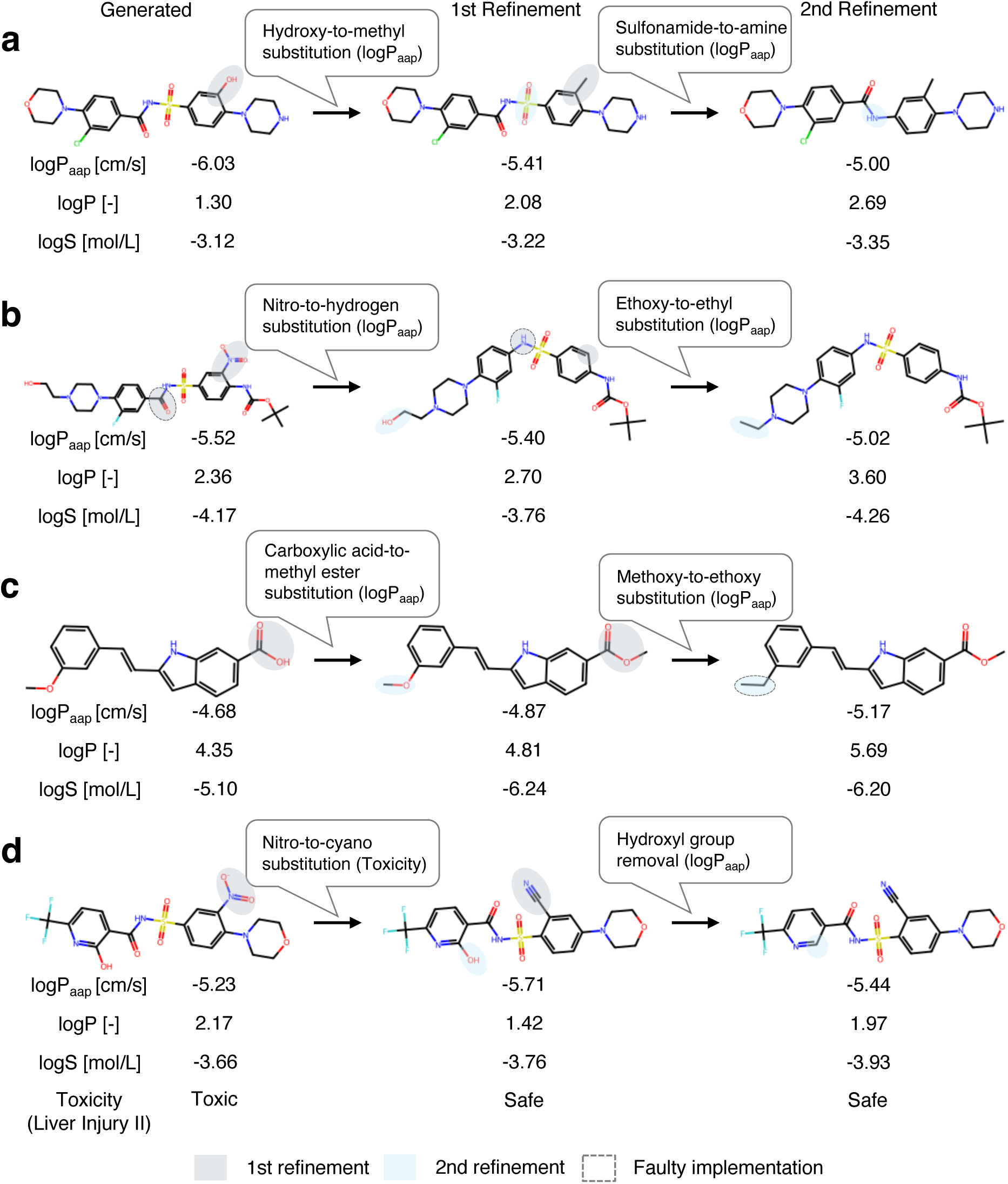
Representative examples of property-aware molecular refinement. Each case shows structural modifications aimed at improving a specific target property (indicated in parentheses), as proposed by the agent’s reasoning output. the Rule of Three, and Oprea’s lead-likeness filter. ^46–50^ Figure 7d shows a steady rise in the number of molecules passing four or more of these rules, increasing from 29 to 44 and then to 52, indicating improved developability profiles after refinement.

However, refinements are not always successful or precise. Two common failure modes are observed: (i) unintended SMILES modifications that diverge from the agent’s stated intent, and (ii) correctly executed modifications that fail to improve the target property. For case (i), in Figure 6b, the agent intended a nitro-to-hydrogen substitution and successfully applied it, but also unintentionally removed a carbonyl group; nonetheless, the predicted permeability improved. Additionally, in Figure 6c, an intended methoxy-to-ethoxy substitution was improperly implemented, resulting in the loss of an oxygen atom and decreased permeability. For case (ii), even when the modification is applied correctly, the desired outcome may not be achieved, as seen in the first-round refinement of Figure 6c, where the permeability worsened despite the intended structural change.

Quantitatively, in the first refinement round, 57 out of the initial 100 molecules were identified as having low permeability. Among these, approximately 44% showed improved log *P*_app_ values after refinement, 26% declined, and 30% remained unchanged. In the second round, 52% improved, 17% declined, and 31% were unchanged. Toxicity improvements were relatively more limited: 24% and 20% of high-toxicity molecules showed improvement in the first and second rounds, respectively, with most remaining toxic despite modification.

Despite these local failures, the overall impact of refinement is positive across the molecule pool. This is supported by increased QED (Quantitative Estimate of Drug-likeness) scores, which reflect overall drug-likeness based on empirical physicochemical properties.^45^ As shown in Figure 7c, the distribution of high-QED molecules expanded with each refinement iteration: the number of molecules with QED *>* 0.6 increased from 34 in the original set to 49 after the first update, and to 55 after the second. In addition, more molecules satisfied empirical drug-likeness criteria - including Lipinski’s Rule of Five, Veber’s rule, Ghose’s rule,

**Figure 7:**
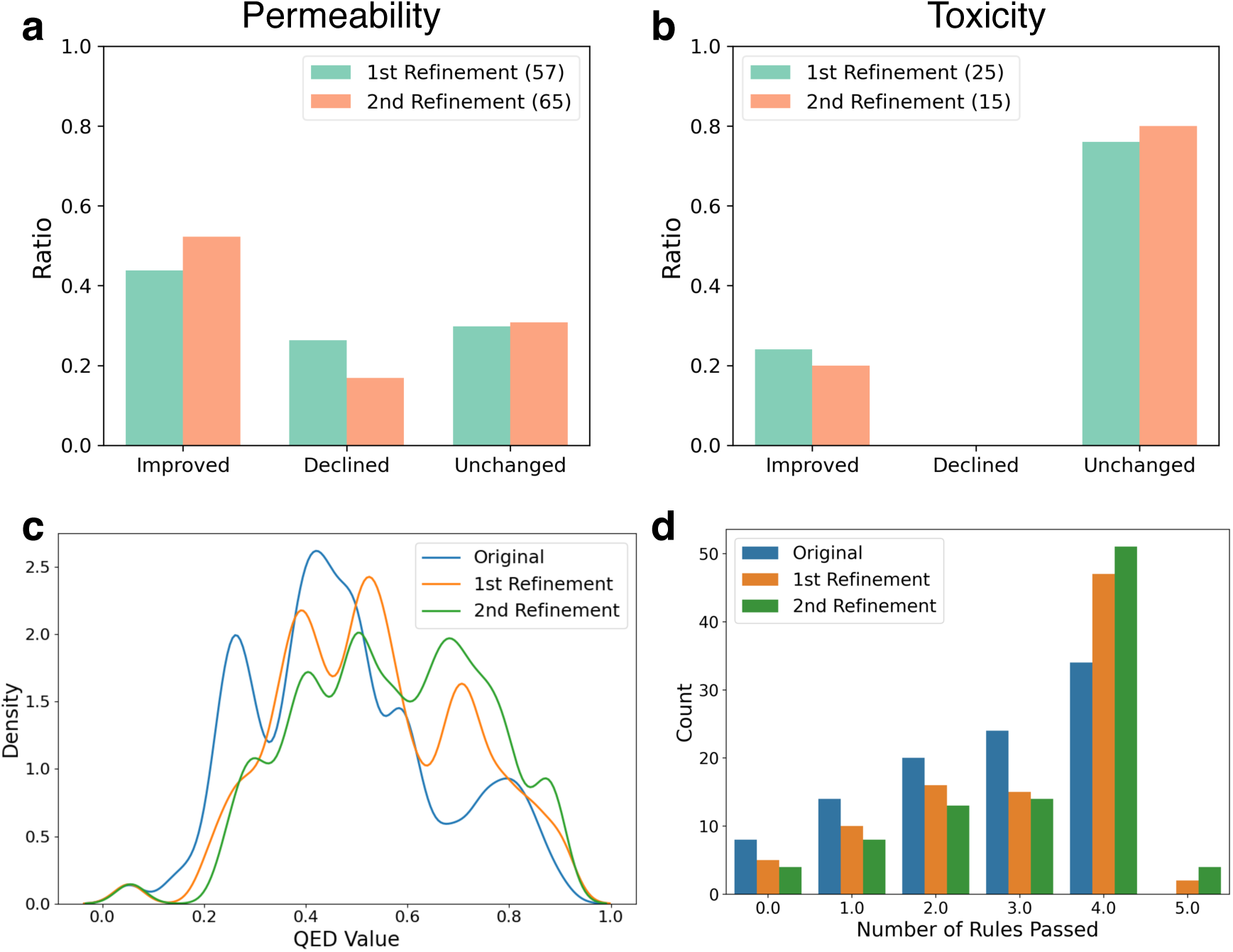
Property changes across iterations of property-aware molecular refinement. **a** and **b** show the proportion of molecules with improved, declined, or unchanged permeability (logP_app_) and toxicity-related properties. The number of molecules targeted in each refinement round is indicated in the legend. **c** Kernel density plot showing the distribution shift in QED scores across refinement iterations. **d** Histogram illustrating the number of molecules satisfying empirical drug-likeness rules; detailed rule definitions are provided in the Methods section.

### 3D Structure Generation

After refining the seed molecule pool through property-aware modifications, we apply empirical filtering criteria to identify high-potential candidates for 3D structural evaluation. Selection is guided by widely accepted drug-likeness rules, including Lipinski’s Rule of Five, Veber’s rule, Ghose’s rule, the Rule of Three, and Oprea’s lead-likeness filter, as detailed in the Table S3. ^46–50^ It is worth noting that this filtering strategy is illustrative, intended to showcase the agent’s generation capabilities. In our workflow, molecules are shortlisted if they satisfy at least three of these five rules, have a QED score above 0.55, and a predicted pKd value greater than 6.0. It is important to note that this criterion is illustrative and intended to demonstrate the pipeline’s capability for structure generation, and may differ from criteria used in specific therapeutic applications. The molecule shown in Figure 8 meets four of the five drug-likeness rules, has a QED score of 0.68, and a predicted pKd of 6.18, making it a candidate for further analysis. Additional examples are provided in Figure S1.

**Figure 8:**
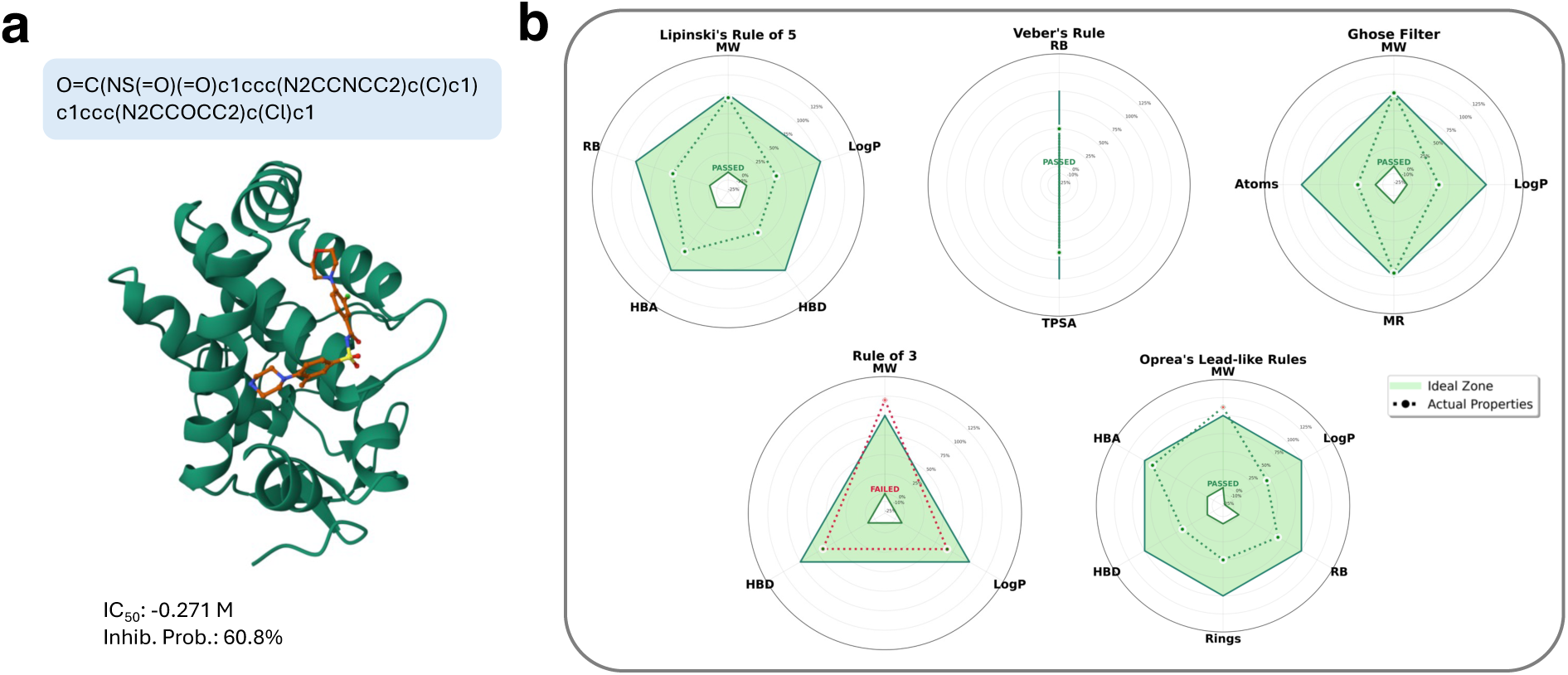
Protein–ligand structure generation and evaluation. **a**, A representative 3D complex predicted by Boltz-2, with estimated IC_50_ and inhibitor probability. **b**, Radar plot showing empirical rule-based evaluation. Green dotted lines indicate the thresholds for each rule, while the purple solid line represents the properties of the selected molecule. See Methods and Table S3 for property definitions.

Once candidates are selected, AgentD uses Boltz-2 to generate 3D protein–ligand complex structures using the ligand’s SMILES and the target protein’s FASTA sequence. These structures provide a foundation for more rigorous downstream evaluations such as molecular docking, MD simulations, and free energy perturbation analyses, to assess binding stability, conformational flexibility, and interaction specificity under biologically relevant conditions. In addition to structural outputs, Boltz-2 returns estimated binding metrics, including IC_50_ values and inhibitor probabilities. In the example shown, the predicted IC_50_ and inhibitor probability indicate moderate (not strong) binding affinity, suggesting that further molecular optimization or experimental validation may be necessary.

## Discussion

The agent reliably retrieves protein and ligand data from both structured databases and web sources. While our current implementation focuses on targeted extraction, future extensions could incorporate large-scale literature mining and relevance scoring to support broader knowledge synthesis. For domain-specific question answering, AgentD’s RAG responses consistently outperform standard LLM outputs in contextual accuracy and mechanistic detail. However, since RAG performance strongly depends on the quality of retrieved literature, extending access beyond open-access sources could further improve its effectiveness.

For molecule generation, AgentD integrates external models such as REINVENT and Mol2Mol using natural language–driven configuration updates, enabling seamless integration of new tools with minimal customization. By leveraging a language model to interpret and modify configuration files based on high-level instructions, the system can flexibly switch between different generative models or sampling strategies without manual code changes. This approach removes the need for model-specific hardcoded logic and allows for rapid adaptation to evolving architectures and file structures. The modular design of AgentD further ensures its long-term adaptability as generative modeling techniques continue to advance.

The property-aware molecular refinement module shows both promise and limitations. AgentD successfully improves target properties such as permeability or toxicity using only its built-in reasoning and domain knowledge. However, molecular optimization remains intrinsically complex due to interdependencies among properties. For example, in Figure 6b, improvements in permeability (logP_app_) lead to declines in lipophilicity (logP) and solubility (logS). Similarly, the first-round refinement in Figure 6c mitigates predicted liver toxicity but worsens permeability. These trade-offs highlight the fundamental challenge of multiproperty optimization in drug design. Despite these complexities, AgentD achieves a net improvement in overall drug-likeness across the molecule pool, as shown in Figure 7c and d. These findings highlight the importance of incorporating multi-objective optimization in future iterations to better balance competing pharmacological goals.

The structure generation task complements the pooling and refinement process by enabling 3D protein-ligand complex generation, which provides critical input for structure-based studies. Although we use general drug-likeness criteria such as Lipinski’s Rule of Five and QED thresholds for demonstration purposes, target-specific filters can be readily implemented in practice. Boltz-2 captures meaningful trends in ligand binding behavior, but its IC_50_ and inhibitor probability predictions should be interpreted as rough approximations that are useful for initial prioritization rather than as substitutes for detailed structure-based evaluation.

## Conclusion

In this study, we introduced AgentD, a modular, LLM-powered agent framework for automating and streamlining key stages of the drug discovery pipeline. By integrating language model reasoning with domain-specific tools and databases, AgentD can: (i) retrieve relevant protein and compound data from structured databases and web sources; (ii) answer domain-specific scientific questions grounded in literature; (iii) generate diverse, context-aware molecules using both de novo and conditional models; (iv) predict pharmacologically relevant properties; (v) refine molecular representations through iterative, property-aware optimization; and (vi) construct protein–ligand complex structures for downstream simulations.

Overall, AgentD marks a step toward general-purpose, AI-driven scientific agents for therapeutic discovery. Its modular architecture supports seamless integration of new models and tools, ensuring continued adaptability as technologies evolve. Future extensions may enhance its capabilities through autonomous molecular dynamics simulations for structural validation, multi-objective molecular optimization, and the generation of candidates conditioned on specific pharmacological profiles.

## Methods

### Large Language Model Agent

Our framework employs OpenAI’s GPT-4o model^51^ as the primary language model for all agentD modules. GPT-4o is an optimized version of GPT-4, which belongs to the Generative Pretrained Transformer family.^52^ The model uses transformer architecture with self-attention mechanisms^53^ to effectively process contextual relationships in text sequences. GPT-4o maintains the strong language understanding capabilities of GPT-4 while offering improved computational efficiency through performance optimizations. We selected GPT-4o specifically for its proven effectiveness in scientific reasoning tasks and its extensive training on chemical and biological literature. We implement the agent framework using LangChain,^54^ a Python library for developing large language model applications. LangChain simplifies the management of complex workflows by handling prompt coordination, response processing, and integration with external tools and APIs.

### Database

To facilitate target-specific molecular design, we utilize two established bioinformatics resources, UniProt and ChEMBL, to access protein sequence data and small molecule representations, respectively. UniProt provides high-quality annotated protein sequences between species, enabling access to functionally characterized targets of therapeutic relevance.^34^ For each therapeutically relevant protein target (e.g., *EGFR* or *TP53*), the agent queries

UniProt’s REST API, restricted to the *Homo sapiens* taxonomy (organism ID: 9606) to retrieve the corresponding amino acid sequence in FASTA format. When existing drugs are available for the target, we subsequently query ChEMBL to obtain SMILES representations of these known compounds.^35^ This ChEMBL query step is only executed if existing drugs for the target are identified through prior web searches; otherwise, this step is skipped.

### ADMET Properties

ADMET properties encompass Absorption, Distribution, Metabolism, Excretion, and Toxicity characteristics that are critical for drug development.^55,56^ These pharmacologically relevant descriptors determine how a compound is absorbed into the bloodstream, distributed across tissues, metabolized by enzymatic systems, eliminated from the body, and whether it poses potential toxic effects. Early prediction of ADMET properties is essential for prioritizing compounds with favorable biopharmaceutical and safety profiles, thereby reducing downstream attrition during drug development.^57,58^

As part of this multi-stage assessment, we integrated Deep-PK, an ADMET prediction framework that operates on SMILES input and internally employs a Message Passing Neural Network (MPNN) to capture the atomic and topological features of each molecule.^39^ This architecture allows for graph-based encoding of chemical structures by propagating information across atom–bond interactions. Candidate ligands, whether recovered from ChEMBL or generated, are submitted to the Deep-PK REST API via their SMILES strings, and the resulting ADMET profiles are parsed to prioritize compounds with favorable pharmacokinetic and safety attributes. The complete set of predicted properties was summarized in Supplementary Information Table S2.

### Binding Affinity

Binding affinity to the biological target is a fundamental determinant of therapeutic potential, serving as a key predictor of drug potency and selectivity. We employ two complementary strategies for affinity prediction. In the property prediction task, the sequence-based BAPULM model employs a dual encoder architecture to estimate the dissociation constant (*K_d_*) from protein amino acid sequences and ligand SMILES representations. It integrates two domain-specific pre-trained language models: ProtT5-XL-U50 for proteins and Mol-Former for small molecules.^59,60^ Each encoder generates latent embeddings tailored to its respective input, which are then projected into a shared latent space using learnable feedforward projection heads. Subsequently, a predictive head processes these joint representations to estimate the binding affinity, reported as *pK_d_* = − log_10_(*K_d_*).^40^ Given its computational efficiency and reliance solely on sequence-level inputs, it is well-suited for early-stage screening of large ligand libraries based on predicted binding affinity.

In the structure generation task, the structure-based model Boltz begins with the same sequence and SMILES input but internally generates 3D protein–ligand complex structures. These conformations are then used to predict the half-maximum inhibitory concentration (IC_50_), reflecting the inhibitory efficacy of a compound in biochemical assays.^61^ In addition to regression-based affinity values, Boltz also outputs inhibitor probability scores, indicating the likelihood that a given ligand acts as an active binder. Although *K_d_* and IC_50_ originate from different experimental setups, they are related through the Cheng-Prusoff equation.^62^

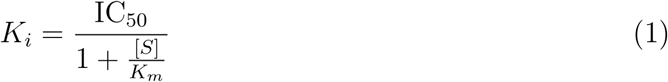

where *K_i_* approximates *K_d_* under certain biochemical conditions.

### SMILES Generation

We utilize two molecular generators: REINVENT for de novo molecular design and Mol2Mol for molecular optimization.^36–38^ Both generators utilize recurrent neural networks and transformer architectures and are embedded within machine learning optimization algorithms, including reinforcement learning and transfer learning.^37,63^

REINVENT performs de novo molecular generation using sequence-based models that capture SMILES token probabilities in an autoregressive manner. The models are trained via teacher-forcing on large SMILES datasets to learn chemical syntax and generate valid molecules.^64^ Reinforcement learning optimization employs a policy gradient scheme using the “Difference between Augmented and Posterior” (DAP) strategy, where scalar scores are computed for generated molecules based on user-defined scoring functions.^65^ An augmented likelihood combines this reward signal with a fixed prior model likelihood, providing regularization to maintain chemical plausibility while optimizing toward desired properties.

Mol2Mol performs conditional generation, accepting input SMILES strings and generating structurally similar molecules within user-defined similarity constraints.^38^ The transformer-based model was trained on over 200 billion molecular pairs from PubChem with Tanimoto similarity ≥ 0.50. Training employed ranking loss to directly link negative log-likelihood to molecular similarity. The model supports multinomial sampling with temperature control and beam search decoding, enabling controlled exploration around known compounds while preserving structural relationships to the input molecule.

### Protein-Ligand Complex Structure

Boltz-2 is a structure-based deep learning model designed to jointly predict 3D protein–ligand complex structures by integrating protein folding and ligand binding into a unified framework.^42^ The model takes as input a protein FASTA sequence and a ligand SMILES string, and simultaneously infers the full atomic conformation of the protein as well as the bound pose of the ligand within the predicted binding pocket. Unlike traditional docking pipelines that require experimentally resolved protein structures, Boltz-2 performs *ab initio* structure prediction, enabling end-to-end modeling from sequence alone.

Its architecture builds upon the Evoformer ^17^ stack and SE(3)-equivariant transformer modules, incorporating interleaved attention mechanisms to capture long-range dependencies both within the protein sequence and between the protein and ligand.^41,42,53^ This allows Boltz-2 to reason over flexible ligand conformations and protein-side chain rearrangements in a physics-aware manner. By directly predicting all-atom 3D coordinates, the model enables rapid generation of realistic protein–ligand complexes suitable for downstream scoring.

### Evaluation Metrics

To evaluate the drug-likeness and pharmacokinetic relevance of the generated molecules, we applied five widely used rule-based filters: Lipinski’s Rule of Five,^46^ Veber’s Rule, ^47^ Ghose Filter,^48^ Rule of Three (Ro3), ^49^ and Oprea’s Lead-like Rule.^50^ The specific numerical criteria for each rule are summarized in Table S3 of the Supporting Information. Molecular descriptors were computed using RDKit where necessary.

Lipinski’s Rule of Five includes five criteria: molecular weight (MW) ≤ 500 Da, LogP ≤ 5, hydrogen bond donors (HBD) ≤ 5, hydrogen bond acceptors (HBA) ≤ 10, and rotatable bonds ≤ 10. Although the original rule included only four properties, the rotatable bond constraint is now widely adopted to better capture oral bioavailability. A molecule was considered compliant if it satisfied at least four of the five conditions.

Veber’s rule assesses polarity and flexibility using two thresholds: topological polar surface area (TPSA) ≤ 140 Å^2^ and rotatable bonds ≤ 10. Both conditions must be met for compliance.

The Ghose filter identifies well-balanced drug-like molecules based on MW between 160–480 Da, LogP between –0.4 and 5.6, molar refractivity (MR) between 40–130, and heavy atom count between 20–70.

The Rule of Three, designed for fragment-based discovery, applies stricter limits: MW *<* 300 Da, LogP ≤ 3, and HBD ≤ 3. It is used to identify small, synthetically accessible fragments with potential for optimization.

Oprea’s Lead-like Rule defines criteria suitable for early-stage lead optimization: MW between 200–450 Da, LogP between –1 and 4.5, HBD ≤ 5, HBA ≤ 8, rotatable bonds ≤ 8, and aromatic rings ≤ 4. One violation among these conditions was permitted to allow flexibility in lead selection.

### Technology Use Disclosure

We used ChatGPT and Claude to help with grammar and typographical corrections during the preparation of this preprint manuscript. The authors have carefully reviewed, verified, and approved all content to ensure accuracy and integrity.

## Code Availability Statement

The code that supports this study’s findings can be found in the following publicly available GitHub repository: https://github.com/hoon-ock/AgentD.

## Acknowledgement

The authors gratefully acknowledge support from the H. Robert Sharbaugh Presidential Fellowship.

## Author declarations

### Author Contributions

J.O., A.B.F. conceived the study, and J.O., R.S.M., A.C., N.S.A., and S.B. conducted the experiments, analyzed the data, and wrote the original draft. R.S.M., N.S.A., S.B., A.C. contributed to experimental design and data analysis. A.B.F. supervised the project, provided funding acquisition, and reviewed the manuscript.

