## Supporting Information for "Large Language Model Agent for Modular Task Execution in Drug Discovery"

#### Contents

**S1 Domain-specific Question Answering** **S2**

**S2 ADMET Properties** **S6**

**S3 Identified Unfavorable Drug Properties** **S11**

**S4 Structure Generation Examples** **S12**

**S5 Drug-likeness Filters** **S13**

**References** **S14**

### S1 Domain-specific Question Answering

We conducted an additional evaluation question that examined AgentD's ability to extrapolate mechanistic findings to broader therapeutic contexts by asking about the implications of combining ABT-199 and proteasome inhibitors for treating solid tumors compared to hematologic malignancies. This question tests the system's capacity to synthesize domain-specific knowledge and provide clinically relevant insights beyond the immediate scope of the reference paper. Figure S1 shows the actual responses from both AgentD with RAG and the standard GPT-4o model to illustrate these differences in clinical reasoning and mechanistic understanding.

We evaluated responses using the same Weller et al. study, which primarily focuses on the mechanistic basis of drug synergy through NOXA upregulation and MCL-1 inhibition.[S1] While this question extends beyond the paper's direct experimental scope, it remains grounded in the mechanistic principles described in the work. Both AgentD with RAG and the standard GPT-4o model were asked to discuss the therapeutic implications for different cancer types.

The RAG-augmented response demonstrates superior clinical reasoning by directly connecting the paper's mechanistic findings to therapeutic applications. AgentD correctly identifies that the combination could "significantly impact the treatment of both solid tumors and hematologic malignancies" through the dual mechanism of direct BCL-2 inhibition and indirect MCL-1 inhibition via NOXA activation. Importantly, the response provides specific examples, noting effectiveness in "solid tumors, such as soft tissue sarcoma" and established efficacy in hematologic malignancies including "chronic lymphocytic leukemia and multiple myeloma."

**Domain-specific question:**
What are the implications of the findings regarding the combinatorial use of ABT-199 and proteasome inhibitors for treating solid tumors compared to hematologic malignancies?

**AgentD (w/ RAG):**

The findings indicate that combining ABT-199 (venetoclax) with proteasome inhibitors (PIs) could significantly impact the treatment of both solid tumors and hematologic malignancies. In solid tumors, such as soft tissue sarcoma, this combination synergistically induces apoptosis by activating the pro-apoptotic protein NOXA, which inhibits the anti-apoptotic protein MCL-1, enhancing ABT-199's efficacy. This dual action—direct BCL-2 inhibition and indirect MCL-1 inhibition—suggests effectiveness in solid tumors overexpressing these proteins. In hematologic malignancies, where ABT-199 is already effective, particularly in chronic lymphocytic leukemia and multiple myeloma, combining it with PIs could improve outcomes by overcoming resistance mechanisms involving MCL-1. These findings support further clinical evaluation of this combination in various cancers, potentially expanding therapeutic options for both solid and hematologic malignancies.

**Standard GPT4o:**

The findings on the combinatorial use of ABT-199 (venetoclax) and proteasome inhibitors have several implications for the treatment of solid tumors compared to hematologic malignancies. Here's a breakdown:

1. Mechanistic Synergy:
Hematologic Malignancies: ABT-199, a BCL-2 inhibitor, is particularly effective because these cancers often rely heavily on the BCL-2 pathway for survival. When combined with proteasome inhibitors, which block protein degradation, the apoptotic pathways are further enhanced, leading to increased cancer cell death.
Solid Tumors: These cancers might not be as dependent on BCL-2 as hematologic malignancies. However, the combination could still disrupt protein homeostasis and induce apoptosis by targeting multiple pathways.

2. Clinical Efficacy:
Hematologic Malignancies: Clinical trials have shown promising results due to the high BCL-2 expression. The combinatorial approach can enhance responses in resistant cases.
Solid Tumors: The efficacy might be more variable due to the diverse nature of these tumors and their reliance on other anti-apoptotic proteins like MCL-1 or BCL-XL.
Resistance Mechanisms:

Hematologic Malignancies: Resistance can develop through mutations or upregulation of alternative survival pathways.
Solid Tumors: The diverse genetic landscape means multiple resistance mechanisms could limit effectiveness.

3. Toxicity and Side Effects:
Combining ABT-199 with proteasome inhibitors can potentially increase toxicity, necessitating careful dose optimization and monitoring in both cancer types.

4. Biomarker Development:
Identifying biomarkers to predict response can help tailor the combination therapy more effectively, particularly in solid tumors where response rates might be heterogeneous.

5. Future Research Directions:
Continued research is needed to understand the specific tumor types and settings where this combination is most beneficial, and to develop strategies to overcome resistance in solid tumors.
Overall, while the combination has shown efficacy in hematologic cancers, its application in solid tumors requires a nuanced understanding of tumor biology and careful clinical evaluation.

Figure S1: AgentD response demonstrating domain-specific question answering capabilities. The question extrapolates beyond the direct scope of the source article.

The RAG response captures the key therapeutic insight that this combination approach could "overcome resistance mechanisms involving MCL-1," which aligns closely with the ground truth explanation that the strategy "could potentially overcome intrinsic resistance mechanisms in solid tumors that are less dependent on BCL-2 but still rely on MCL-1 for survival." This mechanistic understanding is crucial for clinical translation, as it identifies the specific molecular rationale for why the combination might succeed where single-agent BCL-2 inhibition fails.

In contrast, the standard GPT-4o response, while comprehensive and well-structured, provides a generic framework that could apply to virtually any combination therapy discussion. The response covers broad categories such as "mechanistic synergy," "clinical efficacy," and "resistance mechanisms" but fails to reference the specific NOXA-mediated pathway that makes this particular combination therapeutically promising. Critical omissions include the ISR activation, ATF3/ATF4 transcriptional regulation, and the specific role of MCL-1 inhibition in overcoming resistance.

The standard response does acknowledge that "solid tumors might not be as dependent on BCL-2 as hematologic malignancies" and mentions reliance on "other anti-apoptotic proteins like MCL-1 or BCL-XL," but it lacks the mechanistic foundation to explain how the combination specifically addresses these dependencies. This represents a missed opportunity to provide actionable clinical insights based on the underlying biology.

Furthermore, while the standard response discusses general considerations such as toxicity management and biomarker development, it does so without the mechanistic context that would guide these clinical decisions. The RAG response, by contrast, grounds its recommendations in the specific findings about NOXA upregulation and MCL-1 inhibition, providing a more targeted foundation for clinical development.

This evaluation demonstrates that even for questions that extend beyond the immediate experimental scope of the reference literature, RAG-augmented responses maintain superior clinical relevance by preserving the mechanistic foundation that underlies therapeutic potential. The ability to connect specific molecular mechanisms to broader therapeutic applications represents a critical advantage for biomedical question answering systems, particularly in translational research contexts where mechanistic understanding directly informs clinical strategy.

#### S2 ADMET Properties

Table S1: Complete list of ADMET properties predicted for all compounds in this study.

| Category | Property | Description |
|---|---|---|
| **Absorption** | Caco-2 Permeability (log-Paap) | Permeability across Caco-2 cell monolayers |
| | Human Intestinal Absorption | Fraction absorbed in human intestine |
| | Human Oral Bioavailability (20%) | Probability of achieving >20% oral bioavailability |
| | Human Oral Bioavailability (50%) | Probability of achieving >50% oral bioavailability |
| | MDCK Permeability | Madin-Darby Canine Kidney cell permeability |
| | P-Glycoprotein Inhibitor | Inhibition of P-glycoprotein efflux pump |
| | P-Glycoprotein Substrate | Substrate of P-glycoprotein efflux pump |
| | Skin Permeability | Dermal absorption coefficient |
| **Distribution** | Blood-Brain Barrier Penetration | CNS penetration capability |
| | Blood-Brain Barrier (log BB) | Blood-brain barrier partition coefficient |
| | Fraction Unbound (Human) | Unbound fraction in human plasma |

*Continued on next page*

Table S1 – *Continued from previous page*

| Category | Property | Description |
|---|---|---|
| | Plasma Protein Binding | Extent of protein binding in plasma |
| | Volume of Distribution | Steady-state volume of distribution |
| **Metabolism** | BCRP Substrate | Breast Cancer Resistance Protein substrate |
| | CYP1A2 Inhibitor | Cytochrome P450 1A2 inhibition |
| | CYP1A2 Substrate | Cytochrome P450 1A2 substrate |
| | CYP2C19 Inhibitor | Cytochrome P450 2C19 inhibition |
| | CYP2C19 Substrate | Cytochrome P450 2C19 substrate |
| | CYP2C9 Inhibitor | Cytochrome P450 2C9 inhibition |
| | CYP2C9 Substrate | Cytochrome P450 2C9 substrate |
| | CYP2D6 Inhibitor | Cytochrome P450 2D6 inhibition |
| | CYP2D6 Substrate | Cytochrome P450 2D6 substrate |
| | CYP3A4 Inhibitor | Cytochrome P450 3A4 inhibition |
| | CYP3A4 Substrate | Cytochrome P450 3A4 substrate |
| | OATP1B1 Substrate | Organic Anion Transporting Polypeptide 1B1 |
| | OATP1B3 Substrate | Organic Anion Transporting Polypeptide 1B3 |
| **Excretion** | Clearance | Total body clearance |
| | Half-Life | Elimination half-life |
| | OCT2 Substrate | Organic Cation Transporter 2 substrate |
| **Toxicity** | AMES Mutagenicity | Bacterial mutagenicity test |

*Continued on next page*

Table S1 – *Continued from previous page*

| Category | Property | Description |
|---|---|---|
| | Avian Toxicity | Acute toxicity to birds |
| | Bee Toxicity | Acute toxicity to honeybees |
| | Bioconcentration Factor | Bioaccumulation potential |
| | Biodegradation | Environmental biodegradability |
| | Carcinogenicity | Carcinogenic potential |
| | Crustacean Toxicity | Acute toxicity to crustaceans |
| | Daphnia Toxicity | Acute toxicity to *Daphnia magna* |
| | Eye Corrosion | Severe eye damage potential |
| | Eye Irritation | Eye irritation potential |
| | Fathead Minnow Toxicity | Acute toxicity to *Pimephales promelas* |
| | Hepatotoxicity (DILI) | Drug-induced liver injury |
| | Hepatotoxicity (Alternative) | Alternative liver injury prediction |
| | Maximum Tolerated Dose | Highest non-toxic dose |
| | Micronucleus Test | Chromosomal damage potential |
| | Nuclear Receptor AhR | Aryl hydrocarbon receptor activation |
| | Nuclear Receptor AR | Androgen receptor binding |
| | Nuclear Receptor AR-LBD | Androgen receptor ligand binding domain |
| | Nuclear Receptor Aromatase | Aromatase enzyme inhibition |
| | Nuclear Receptor ER | Estrogen receptor binding |

*Continued on next page*

Table S1 – *Continued from previous page*

| **Category** | **Property** | **Description** |
|---|---|---|
| | Nuclear Receptor ER-LBD | Estrogen receptor ligand binding domain |
| | Nuclear Receptor GR | Glucocorticoid receptor binding |
| | Nuclear Receptor PPAR-$\gamma$ | Peroxisome proliferator-activated receptor $\gamma$ |
| | Nuclear Receptor TR | Thyroid receptor binding |
| | Rat Acute Toxicity | Acute oral toxicity in rats |
| | Rat Chronic Toxicity | Chronic oral toxicity in rats |
| | Respiratory Toxicity | Respiratory system toxicity |
| | Skin Sensitization | Dermal sensitization potential |
| | Stress Response ARE | Antioxidant response element activation |
| | Stress Response ATAD5 | ATAD5 genotoxicity pathway |
| | Stress Response HSE | Heat shock response element |
| | Stress Response MMP | Mitochondrial membrane potential |
| | Stress Response p53 | p53 tumor suppressor pathway |
| | *Tetrahymena pyriformis* | Acute toxicity to *T. pyriformis* |
| | hERG Inhibition | Human ether-à-go-go-related gene K+ channel |
| **General** | Boiling Point | Boiling point temperature |
| | Hydration Free Energy | Free energy of hydration |
| | log D (pH 7.4) | Distribution coefficient at physiological pH |

*Continued on next page*

Table S1 – *Continued from previous page*

| **Category** | **Property** | **Description** |
|---|---|---|
| | log P | Octanol-water partition coefficient |
| | log S | Aqueous solubility |
| | log Vapor Pressure | Vapor pressure |
| | Melting Point | Melting point temperature |
| | pKa (Acidic) | Acid dissociation constant |
| | pKa (Basic) | Base dissociation constant |
| | pKd (Acidic) | Alternative acid dissociation constant |

### S3 Identified Unfavorable Drug Properties

The molecular optimization process focuses on identifying unfavorable properties and refining SMILES to improve their ADMET profiles. Table S2 summarizes the distribution of properties flagged by AgentD across optimization rounds, grouped by ADMET categories.

An observed limitation is the agent's frequent selection of $\log P_{\mathrm{app}}$ as a risk factor—despite it being just one among 74 possible properties. This suggests a selection bias, potentially due to the ordering of properties in the input dictionary. Since $\log P_{\mathrm{app}}$ appears near the top of the dictionary entry, the agent may disproportionately attend to it during its reasoning process. Future iterations can address this by randomizing property order or introducing attention calibration techniques.

Table S2: Distribution of weakness properties after 1st and 2nd round of SMILES optimization. The weakness property represents the most critical ADMET deficiency targeted for improvement.

| **ADMET Category** | **Specific Property** | **1st Round (%)** | **2nd Round (%)** |
|---|---|---|---|
| Absorption | Caco-2 (logPaap) | 59.6 | 69.5 |
| Toxicity | AMES Mutagenesis | 14.1 | 7.4 |
| | Liver Injury I (DILI) | 5.1 | 4.2 |
| | hERG Blockers | 4.0 | 2.1 |
| | Carcinogenesis | 3.0 | 1.1 |
| General Properties | logP | 5.1 | 3.2 |
| | logS | 1.0 | – |
| Other | Absorption/Human Oral Bioavailability (20% and 50%), Invalid SMILES, and other general entries. | 6.0 | 8.4 |
| **Total Entries** | | **99** | **95** |

### S4 Structure Generation Examples

For the examples shown in Figure S2, we applied stricter rule-based filters and slightly more lenient pKd thresholds compared to those used in the main manuscript. Specifically, molecules were selected if they passed more than three rule-based filters, had a QED score above 0.6, and a predicted pKd greater than 5.5. From the set of molecules that underwent two rounds of refinement, three examples were randomly selected for illustration.

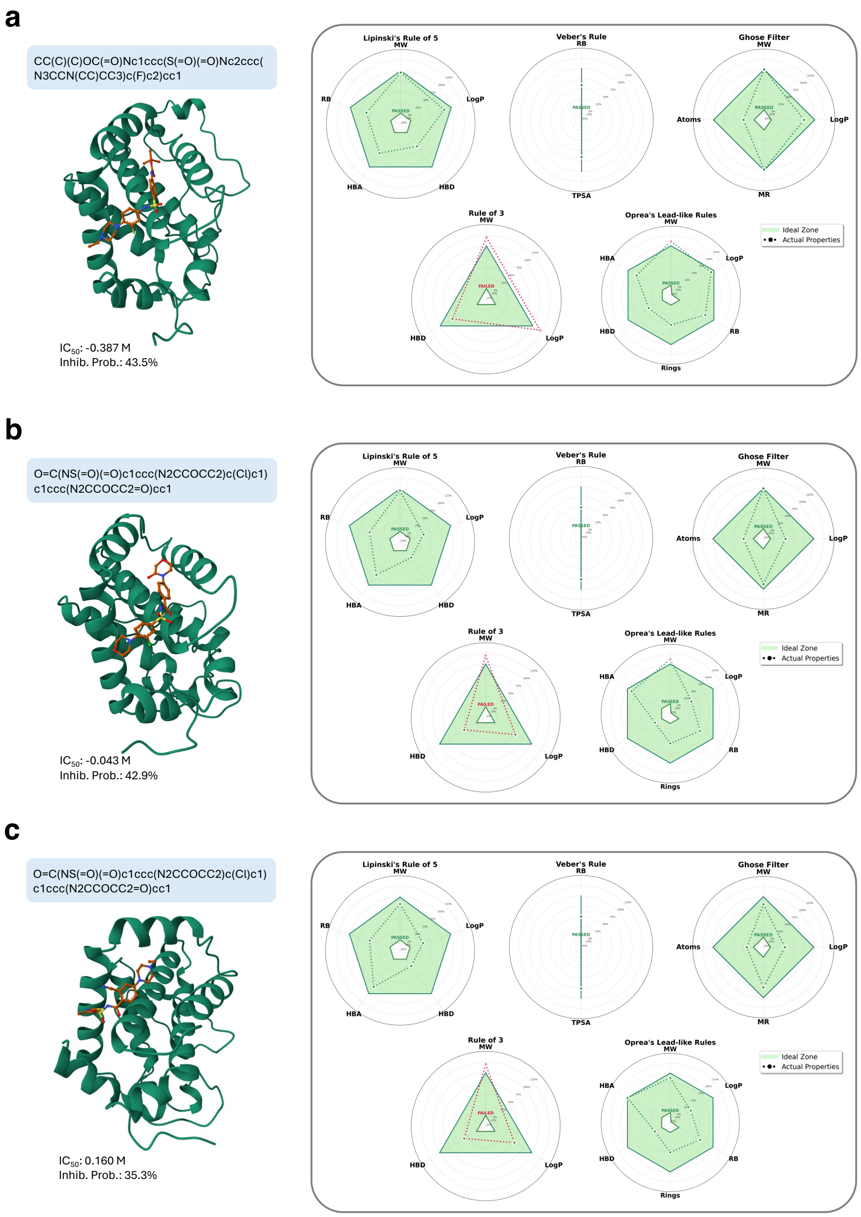


Figure S2: Additional examples of protein–ligand complex generation and evaluation.

### S5 Drug-likeness Filters

The listed properties were calculated using RDKit with molecular SMILES as input. Note that these rules serve as illustrative examples and are not the definitive criteria for selecting candidates for 3D structure generation.

Table S3: Summary of rule-based drug-likeness filters and their physicochemical criteria.

| **Rule** | **Criterion** | **Threshold / Range** | **Pharmacological Rationale** |
|---|---|---|---|
| Lipinski's Rule of Five[S2,S3] | Molecular Weight (MW) | $\leq$ 500 Da | High molecular weight is associated with poor intestinal absorption due to size-related transport limitations. |
| | logP | $\leq 5$ | High lipophilicity (logP $\leq$ 5) correlates with poor aqueous solubility and passive permeability. |
| | H-bond Donors (HBD) | $\leq 5$ | Excessive H-bond donors increase polarity, reducing membrane permeability. |
| | H-bond Acceptors (HBA) | $\leq 10$ | Too many acceptors increase polarity, hindering passive diffusion. |
| | Rotatable Bonds (RB) | $\leq$ 10 rotatable bonds | High flexibility and polar surface area reduce the likelihood of oral activity. |
| Veber Rule[S3] | TPSA | $\leq$ 140 Å$^2$ | High polar surface area decreases oral bioavailability by reducing passive diffusion. |
| | Rotatable Bonds (RB) | $\leq 10$ | Excess flexibility increases entropy, reducing oral bioavailability and metabolic stability. |
| Ghose Filter[S4] | Molecular Weight (MW) | 160–480 Da | Balances molecular size for optimal binding and permeability. |
| | logP | $-0.4$ to 5.6 | Ensures moderate lipophilicity for both solubility and membrane crossing. |
| | Molar Refractivity (MR) | 40–130 | Captures molecular volume and polarizability, influencing receptor binding. |
| | Atom Count | 20–70 | Reflects a size range favorable for drug-likeness and synthetic accessibility. |
| Rule of Three (Ro3)[S5] | Molecular Weight (MW) | $<$ 300 Da | Smaller fragments are preferred in fragment-based drug discovery for lead optimization. |
| | logP | $\leq 3$ | Low lipophilicity promotes solubility in fragment-like compounds. |
| | H-bond Donors (HBD) | $\leq 3$ | Reduces polarity, supporting fragment permeability and binding. |
| Oprea Lead-like Filter[S6] | Molecular Weight (MW) | 200–450 Da | Ideal size range for optimization into drug-like compounds. |
| | logP | $-1$ to 4.5 | Moderate hydrophobicity for balanced solubility and permeability. |
| | Rotatable Bonds (RB) | $\leq 8$ | Lower conformational entropy improves binding efficiency. |
| | Aromatic Rings | $\leq 4$ | Limits excessive aromaticity, which can affect solubility and toxicity. |
| | H-bond Donors (HBD) | $\leq 5$ | Controls molecular polarity and improves membrane permeability. |
| | H-bond Acceptors (HBA) | $\leq 8$ | Keeps polarity within bounds for favorable pharmacokinetics. |
